# The structure of NAD^+^ consuming protein *Acinetobacter baumannii* TIR domain shows unique kinetics and conformations

**DOI:** 10.1101/2023.05.19.541320

**Authors:** Erik Klontz, Juliet O. Obi, Yajing Wang, Gabrielle Glendening, Jahid Carr, Constantine Tsibouris, Sahthi Buddula, Shreeram Nallar, Alexei S. Soares, Dorothy Beckett, Jasmina S. Redzic, Elan Eisenmesser, Cheyenne Palm, Katrina Schmidt, Alexis H. Scudder, Trinity Obiorah, Kow Essuman, Jeffrey Milbrandt, Aaron Diantonio, Krishanu Ray, Daniel Deredge, M LD. Snyder, Greg A. Snyder

**Affiliations:** Division of Vaccine Research, Institute of Human Virology, University of Maryland, School of Medicine, Baltimore, Maryland 21201 USA; Department of Pharmaceutical Sciences, School of Pharmacy, University of Maryland, Baltimore, MD 21201, USA; China Pharmaceutical University, Nanjing, P.R. China; Department of Microbiology and Immunology, University of Maryland, School of Medicine, Baltimore, Maryland 21201 USA; Department of Biochemistry and Molecular Biology at the University of Maryland, School of Medicine, Baltimore, Maryland 21201, USA; Brookhaven National Laboratory, NSLS-II Bldg. 745, Upton, New York 11973, USA; Department of Chemistry and Biochemistry, University of Maryland, College Park, Maryland 20742, USA; Department of Biochemistry and Molecular Genetics, School of Medicine, University of Colorado Denver, School of Medicine, Aurora, CO 80045 USA; Department of Biological Sciences, Towson University, 8000 York Rd, Towson, Maryland 21252 USA; Department of Developmental Biology, Washington University School of Medicine, St. Louis, Missouri 63110, USA

**Keywords:** Toll/Interleukin-1 receptor (TIR), nicotinamide adenine dinucleotide (NAD), hydrolase, bacterial pathogenesis, innate immunity

## Abstract

Toll-like and Interleukin-1/18 receptor resistance (TIR) domain-containing proteins function as important signaling and immune regulatory molecules. TIR domain-containing proteins identified in eukaryotic and prokaryotic species also exhibit NAD+ hydrolase activity in select bacteria, plants, and mammalian cells. We report the crystal structure of the *Acinetobacter baumannii* TIR domain protein (AbTir-TIR) with confirmed NAD^+^ hydrolysis and map the conformational effects of its interaction with NAD^+^ using HDX-MS. NAD^+^ results in mild decreases in deuterium uptake at the dimeric interface. In addition, AbTir-TIR exhibits EX1 kinetics indicative of large cooperative conformational changes which are slowed down upon substrate binding. Additionally, we have developed label-free imaging using 2pFLIM which shows differences in bacteria expressing native and mutant NAD+ hydrolase-inactivated AbTir-TIR^EA^ protein. Our observations are consistent with substrate-induced conformational changes reported in other TIR model systems with NAD+ hydrolase activity. These studies provide further insight into bacterial TIR protein mechanisms and their varying roles in biology.

## Introduction

Bacterial Toll-like and Interleukin-1/18 receptor *(TIR)* resistance proteins have been identified based on sequence motifs conserved with human TIR proteins [1, 2]. Uropathogenic *E. coli* CFT073 and *Brucella Spp.* express soluble TIR domain-containing proteins that act as virulence factors by physically interacting with and disrupting host TIR domain-containing signaling complexes to promote pathogenicity and modulate immunity [3–6]. TIR domain-containing proteins also have been identified both in several additional disease-causing bacteria as well as in non-pathogenic and commensal bacteria, raising questions regarding potential roles they might have apart from their originally described roles in pathogenicity and modulation of mammalian TLR signaling [2, 3, 7–10].

In a recent discovery, TIR domain-containing proteins, which are conserved across biology in bacteria, plants, and animals, were found to exhibit enzymatic function in nicotinamide adenine dinucleotide (NAD^+^) hydrolysis [11, 12]. NAD+ is an essential cofactor and critical regulator of cellular and metabolic function [13]. In humans, TIR-mediated NAD+ hydrolase activity is involved in axonal degradation via the TIR domain-containing adaptor protein Sterile alpha and TIR motif-containing protein SARM[14, 15]. In plants and bacteria, TIR domain-containing proteins with NAD+ hydrolase activity are involved in immune protection against pathogens [12, 16, 17]. Several pathogenic bacterial TIR proteins with confirmed roles in virulence exhibit NAD+ hydrolase activity. Characterized bacterial TIR domain-containing proteins include *Staphylococcus aureus* (TirS)[18, 19], *Pseudomonas aeruginosa* (PumA)[20], Uropathogenic *Escherichia coli* CFT073 (UPEC-TcpC) [3, 4, 21], *Brucella melitensis (TcpB)* [4, 22] and *Acinetobacter baumannii (AbTir-TIR)*[11]. In addition, NAD+ hydrolase activity has been identified in non-pathogenic and commensal bacteria [9, 11, 12, 17]. This newly discovered TIR-mediated enzymatic function lends additional insight and opens new questions regarding the function TIR domain-containing protein may play in bacterial-host interactions. To better understand the molecular mechanisms underlying enzymatically active bacterial TIR proteins, we have performed structure-function studies characterizing the *Acinetobacter baumannii* TIR protein (AbTir-TIR).

*Acinetobacter baumannii* is a gram-negative bacteria and a leading cause of hospital-acquired infections. Due to the emergence of multidrug-resistant (MDR) *A. baumannii* strains that are resistant to nearly all antibiotics classes, it is a bacterial pathogen of global concern. Immune responses against the highly prevalent MDR strain *A. baumannii* CN40 include inflammasome and caspase activation via type I IFN signaling responses mediated by the TIR-domain-containing adapter-inducing interferon-β (TRIF)[23]. AbTir-TIR exhibits NAD^+^ hydrolase activity [11]. Upon cleavage of NAD, AbTir-TIR catalyzes the production of a novel cyclic ADP ribose (cADPR) variant. A subset of TIR domain-containing proteins including select archaeal and bacterial species produce novel cADPR, and the potential functions in metabolism and signaling of this variant have yet to be fully characterized [6], To better understand the mechanism of bacterial TIR NAD+ hydrolase activity and function, we pursued molecular studies of AbTir-TIR. Accordingly, we have used X-ray crystallography to determine the atomic structure of AbTir-TIR and have confirmed its enzymatic activity using liquid chromatography, mass spectrometry and colorimetric assays. In addition, we have used 2pFLIM to image live bacteria label-free that express AbTir-TIR and hydrogen-deuterium exchange mass spectrometry (HDX-MS) to map the kinetics and conformational changes induced by the interaction of NAD+ with AbTir-TIR.

Understanding bacterial TIR protein-mediated regulation of host innate immune responses in the context of NAD+ hydrolase activity could lead to novel antimicrobial strategies for limiting infection, controlling inflammation and modulating immunity and cellular processes.

## Results

### *Acinetobacter baumannii* TIR protein is conserved among bacteria and humans

A GenBank blastP search of the Protein Database (PDB) using *Acinetobacter baumannii* Tir-TIR domain identified several other bacterial TIR domain containing proteins with homology to AbTir-TIR, including structurally characterized TcpB from *Brucella* spp. (PDB ID 4LQC) and PdTIR *Paracoccus denitrificans* PdTIR (PDB 3H16) (sFig. 1). Surprisingly, this search did not identify structures of TIR proteins with NAD^+^ hydrolase activity from plants or animals [12, 16]. A sequence alignment of AbTir-TIR with bacterial TIR proteins with identified NAD+ hydrolase activity shows the conservation of the C-Helix WxxxE motif identified in *Brucella* TcpB to be important for microtubule binding. This motif includes a highly conserved Tryptophan (W) and a Glutamic acid (Glu, E) found at the carboxy terminus that is involved in substrate interactions and necessary for enzymatic activity in most hydrolases [24–26]. This region is highly conserved across plant, bacteria, and mammalian extracellular Toll-like receptor TIR proteins (sFig. 1) [11, 25, 27]. The C-helix WxxxE motif and surrounding region include the NAD^+^ binding site, TIR interactions and autoregulation site in huSARM1 and plant NLR RUN1 [27, 28].

### AbTir-TIR domain is observed to dimerize in solution and in the crystal lattice

The expressed and purified AbTir-TIR domain is observed in solution as a weakly self-associated dimer by size exclusion, analytical ultracentrifugation and within the asymmetric crystal lattice (Fig. 1A-C). TIR domain self-association is consistent with observations for other bacterial TIR protein domains. The size exclusion chromatogram of AbTir-TIR indicated the formation of higher-ordered oligomeric assemblies. The sample separated by SEC exhibits both a single monomer peak and higher ordered species consistent with dimer and soluble aggregate species based on comparison with the size exclusion standard (Fig. 1A). Soluble aggregate and oligomer fractions were confirmed to migrate at a molecular weight similar to monomer peak fractions during SDS-PAGE analysis (data not shown). SEC purified monomer peak fractions were concentrated and screened for crystallization using commercial screens (sFig 2). Small broom-like crystals were identified, reproduced and optimized using streak seeding (sFig 2). Single crystals were isolated and rasterized to identify single crystal diffraction and data was collected using the highly automated microfocus beamline NSLS-II-17-ID AMX [29]. To further confirm the dimer species observed by SEC and in the crystal of AbTir-TIR we performed in solution sedimentation analytical ultracentrifugation (AUC) analysis on the isolated AbTir-TIR monomer SEC peak fraction species [30, 31]. Sedimentation equilibrium acquired on purified AbTir-TIR samples at varying concentrations and speeds indicate weak dimerization with a K_D_ of 1.4μM. Sedimentation velocity performed on concentrated monomer fraction (AbTIR) at 10μM 45K rpm and analyzed using DCDT^+^, indicates a sedimentation coefficient, s_20,w_, of 2.8S, which is consistent with a dimer observed in the crystal (Fig. 1C) [32, 33].

**Figure 1.**
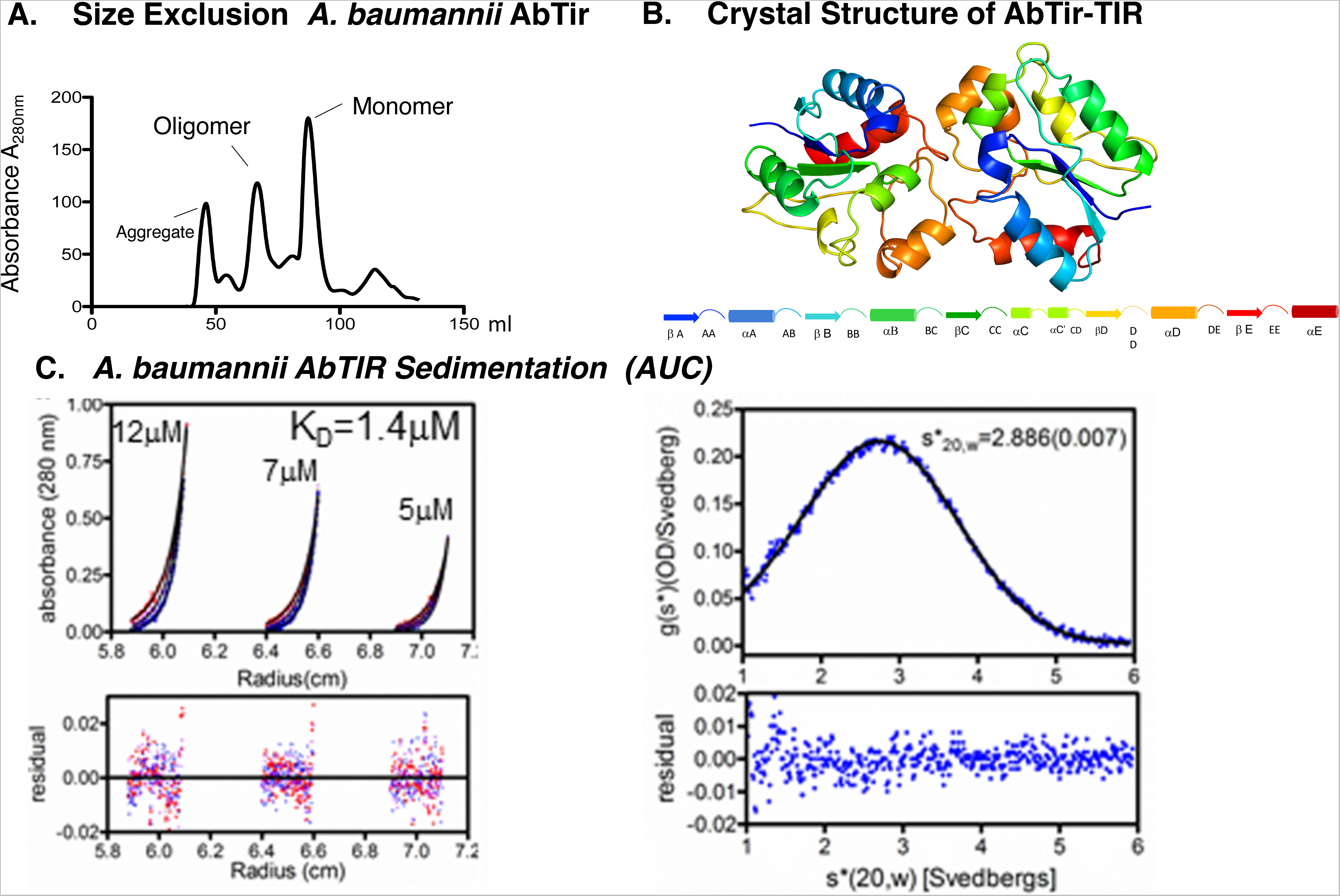
AbTir-TIR is observed as a dimer in solution by SEC, AUC and in the crystal. A. Size exclusion chromatograms showing isolation of stabilized oligomers of full-length AbTir-TIR^E208A^. B. The crystal structure of AbTir-TIR^wt^ domain is observed as a dimer is shown as a colored-coded cartoon ribbon representation. The AbTir-TIR domain shown consists of five alternating alpha helix and beta-strand secondary structures that are gradient colored starting at the amino terminus in blue to the carboxyl-terminal in red (A-E - bottom left to right). C. Analytical ultracentrifugation (AUC) sedimentation profiles of A. baumannii AbTir-TIR. AUC indicates that AbTir-TIR forms a weakly associated dimer in solution. Analysis of sedimentation equilibrium profiles performed at the indicated AbTir-TIR concentrations 12, 7, and 5μM acquired at 23, 26, and 29K rpm indicate dimerization with a K_D_ OF 1.4μM. Sedimentation velocity performed on concentrated monomer fraction (AbTir-TIR) at 10μM 45K rpm and analyzed using DCDT^+^, indicates a sedimentation coefficient, s_20,w_, of 2.8S, consistent with a dimer observed in crystal [32, 33].

### Recombinant AbTir-TIR is enzymatically active

Several assays confirm the enzymatic activity of expressed and purified AbTir-TIR protein. The NAD+ levels of bacteria expressing recombinant AbTir-TIR and the NAD+ hydrolase activity of recombinant purified AbTir-TIR protein were further assessed using the EnzyChrom™ NAD^+^ assay (Fig. 2A-B). AbTir-TIR^wt^ reduces cellular levels of NAD+ upon recombinant expression in *E. coli*. AbTir-TIR^wt^ ^and^ ^E208A^. NAD+ levels in bacterial lysates from cells expressing AbTir-TIR^wt^ and AbTir-TIR^E208A^ were measured 0-6 hr following IPTG induction of AbTIR Tir-TIR^wt^ ^and^ ^E208A^ expression. Expression of catalytically active AbTir-TIR^wt^ promoted significantly greater levels of NAD+ loss in bacterial lysates than did expression of AbTir-TIR^E208A^ (2-way ANOVA, p < 0.0001). For analysis of NAD+ hydrolase activity of recombinant proteins, purified AbTir-TIR^wt^ and AbTir-TIR^E208A^ were incubated with 5 μM of NAD for 10-30 min, and the levels of NAD+ remaining in the samples were measured. Incubation with AbTir-TIR^wt^ resulted in a significant decrease in the levels of NAD+ compared with AbTir-TIR^E208A^ (2-way ANOVA, p<0.0001).

**Figure 2.**
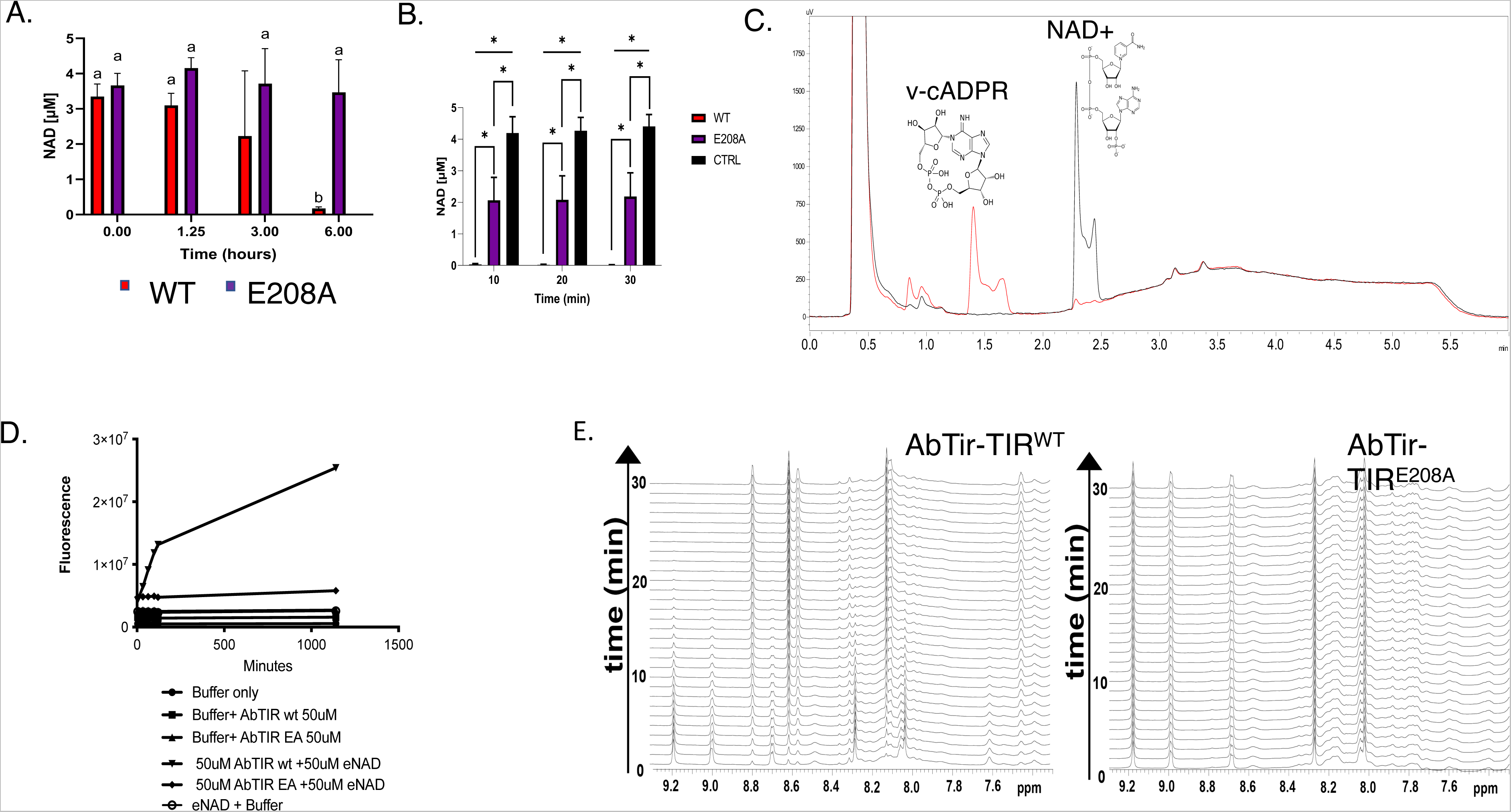
NAD+ Hydrolase activity of AbTir-TIR expressing bacteria and purified recombinant protein. A. Time course of NAD^+^ concentration of bacteria expressing active (AbTir-TIR^wt^) and inactivated (AbTir-TIR^E208A^). B. NAD^+^ concentration overtime of purified recombinant active (AbTir-TIR^wt^), inactivated (AbTir-TIR^E208A^) with buffer control. Asterisk (*) indicates a statistically significant p-value of P ≤ 0.05. C. HPLC trace showing generation of novel cyclic ADP ribose by AbTir-TIR^wt^ protein used in crystallization. HPLC traces of NAD+ protein with AbTir-TIR protein used for crystallization at 0 minutes (black) and after 60min (red) incubation. D. Recombinant AbTir-TIR^wt^ cleaves εNAD. Fluorescence of εNAD in the presence or absence of buffer, AbTir-TIR^wt^, AbTir-TIR^E208A^ measured over time. Buffer only (●), Buffer + AbTir-TIR^wt^ (▪), Buffer + AbTir-TIR^wt^ (▴), εNAD + AbTir-TIR^wt^ (▾), εNAD + AbTir-TIR^208A^ (◆), εNAD + buffer (○). E. Direct visualization of NAD+ catalysis by NMR. One-dimensional NMR spectra of 1 mM NAD+ collected every minute for AbTir-TIR^WT^ (left) and AbTir-TIR^E208A^. Catalysis by AbTir-TIR^WT^ can be observed by the disappearance of the nicotinamide resonances (i.e., 9.2 and 9.0 ppm) that are not observed in the presence of the inactive mutant.

LC-MS trace analysis revealed that recombinant AbTir-TIR^wt^ is capable of hydrolyzing NAD+ into ADPR as well as cyclic ADPR products [11] (Fig. 2C). Additionally, ethano-NAD (εNAD) which yields a fluorescent product upon cleavage was used to assess the enzymatic activity of purified recombinant AbTir-TIR^wt^ in comparison with enzymatic inactivated mutant AbTir-TIR^E208A^ (Fig. 2D) [11, 34]. Only AbTir-TIR^wt^ shows fluorescence of εNAD, indicating NAD+ cleavage in comparison to inactivated mutant AbTir-TIR^E208A^ and buffer controls. One-dimensional NMR spectra also showed NAD+ hydrolase activity of AbTir-AbTIR^WT^ in comparison with inactive mutant AbTir-AbTIR^E208A^ (Fig 2E)[35].

To evaluate AbTir-TIR activity in vivo we developed a label-free assay using the intrinsic fluorescence of NAD(P)H and lifetime (FLT) imaging with 2-photon excitation (2p-FLIM) using live bacteria used for protein expression. Total cellular NAD(P)H fluorescence intensity and average fluorescence lifetimes (FLT) of the same bacteria used for protein expression, for which lysates were assessed for NAD+ levels were measured using 2-pFLIM. Total NAD(P)H and average FLT levels of bacteria expressing enzymatically active (AbTIR^wt/+IPTG^) or inactivated mutant protein (AbTIR^E208A/+IPTG^) along with non-expressing bacteria (AbTIR^wt/-IPTG^) or non-expressing inactivated mutant (AbTIR^E208A/-IPTG^) were quantified (sFig. 3). Differences in both the total NAD(P)H fluorescence intensity and average lifetimes of bacteria-induced to express active AbTir-TIR^wt^ ^(+ITPG)^ or enzymatically inactivated mutant AbTir-TIR^E208A^ ^(+IPTG)^ as well with non-expressing active and inactivated mutant AbTir-TIR^wt/E208A^ ^(-IPTG)^ are observed. In particular, both qualitative and quantitative differences in fluorescence intensities and average lifetimes are observed between expressing and non-expressing active and inactivated mutant AbTir-TIR ^wt/EA^ ^(+ITPG)^ bacteria. Panels representing fluorescence intensities and average lifetimes are quantitated and summarized as histograms (sFig. 3). As expected, differences are observed for bacteria expressing (+IPTG) AbTIR compared with non-expressing (-IPTG), since induction with IPTG is known to affect bacteria growth and metabolism. However, variations in NAD(P)H fluorescence intensities are also observed among active and inactivated bacteria-induced with (+IPTG) to express either AbTir-TIR^wt^ or AbTir-TIR^E208A^ as well as among bacteria not expressing AbTir-TIR^wt^ ^or^ ^E208A^ (sFig. 3 A-D,I). To account for potential differences in the number of bacteria imaged, the average fluorescence intensity for each panel was determined and divided by the total number of bacteria for each panel to yield an average fluorescence intensity per bacteria for each panel. In this analysis, significant differences are observed for inactivated AbTir-TIR^E208A^ (sFig. 3 J). Furthermore, subtracted differences in the average fluorescence lifetimes (FLTs) for NAD(P)H, which is independent of the bacteria concentration and protein expression level, are observed between bacteria expressing either enzymatically active (AbTIR^wt^) or catalytically inactivated mutant AbTir-AbTIR^E208A^ (sFig. 3 E-H and K). Changes in the average fluorescence lifetimes are known to reflect differences in bound and unbound forms of NAD(P)H [36, 37]. Overall, these results indicate AbTir-TIR protein used for structural studies was enzymatically active or inactivated similar to other reports [11, 26].

### The AbTir-TIR crystal structure comparison with other bacterial TIR containing proteins

The structure of *Acinetobacter baumannii* TIR domain (AbTir-TIR) was determined to using X-ray crystallography. The secondary structure architecture is typical of most TIR domains comprised of five alternating beta (β)-strands and alpha (α)-helices (lettered A-E) as initially described [38, 39] (Figure. 1B, sFig 1–2). Conserved loops, helices, individual residues and motifs have been identified and functionally characterized to be important for protein interactions and signaling within several TIR domain-containing proteins and NAD+ hydrolases[40]. The AbTir-TIR is observed as dimeric in solution by SEC and AUC and in the crystal lattice. TIR domain self-association and dimerization, previously shown for SARM TIR hydrolase activity, is consistent with oligomerization of other reported bacterial TIR structures [21, 22, 41–43].

Previously reported bacterial TIR domain-containing structures include *Brucella* TcpB/ BtpA (4LQC,4LZP,4C7M) and *Paracoccus denitrificans* PdTIR (3H16). Most TIR domains are observed as oligomeric species, either dimer and/or tetramer. Individual and TIR domain interactions and loop positions are similar among other reported bacterial TIR domain structures with a root mean square deviation (RMSD) of 1.663 Å (TcpB-4LQC), 1.782 Å (BtpA/TcpB 4LZP) and 2.669 Å (PtDTIR-3H16) for 134, 134, 132, 130 and 137 atom pairs respectively (Fig. 3 and sFig 4). The X-ray structures of AbTir-TIR^wt^ (8G83) and recently reported AbTir-TIR structure (7UWG) are similar [26]. Both apo crystal structures have similar positions for beta strands, alpha helices and intervening loops with an overall RMSD of 0.839 Å. In particular, similar positions for individual residues contained within the WxxxE motif are observed. Both W204 and E208 which are important for selectivity and enzymatic function are observed, solvent exposed and facing outward in the absence of substrate. By contrast, large conformational changes are observed throughout the entirety of the TIR domain structure when comparing apo unliganded X-ray crystal structures (8G83 and 7UWG) with NAD analog bound (3AD) cryoEM structure (AbTir-TIR-3AD 7UXU) Fig 3. Comparing apo X-ray and substrate analog bound cryoEM structures and RMSD of 4.834 is observed (8G83 and 7UXU). Several secondary structures and loops undergo repositioning upon binding substrate analog. These include, conserved residues known to be important for hydrolase activity. For example, both W204 and E208 contained within the WxxxE loop are reoriented inward facing and involved in binding to substrate analog.

**Figure 3.**
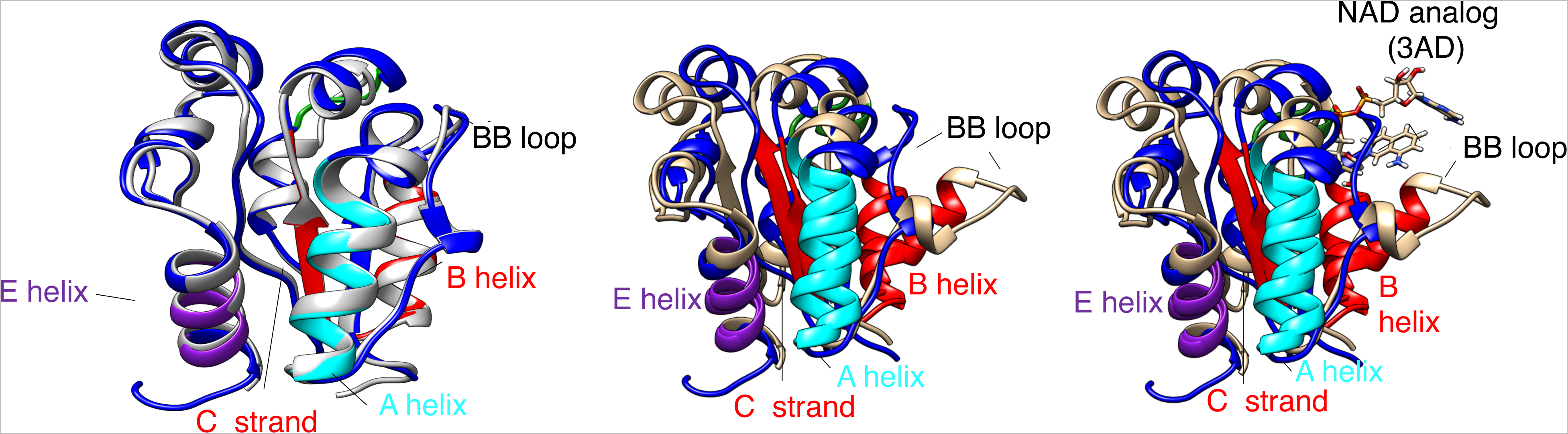
Least squares comparison of unliganded and NAD analog bound X-ray and cryo-EM AbTir-TIR domains. A. Least squares comparison of crystal structures of AbTIr-TIR 8G83(blue) and 7UWG(grey), left panel. Least squares comparison of X-ray AbTir-TIR 8G83 (blue) and cryoEM structure 7UXU (tan) without (middle panel) and with NAD analog 3AD shown (right panel). HDX-MS peptides with changes in EX1 kinetics and conformations upon binding εNAD are colored as shown in Figure 4. A helix (cyan) FVRPLAETLQQL, B helix, BC loop and C beta strand (red) RQKIDSGLRNSKYGTVVL and E helix IAHQLAD (purple).

### AbTir-TIR exhibits unique kinetics and large conformational changes which are slowed down in the presence of substrate

We sought to further characterize AbTir-TIR’s structural dynamics and the conformational consequence of its interaction with NAD+ substrate using hydrogen-deuterium exchange-mass spectrometry (HDX-MS). HDX-MS allows in-solution mapping of interfaces, interactions and allosteric effects by providing comparative peptide-level kinetics of deuterium uptake at global time scales. Sequencing reactions of the AbTir-TIR^wt^ recombinant protein used for crystallization resulted in near complete peptide coverage for AbTir-TIR^wt^ (Fig. S6). The peptide deuteration levels were evaluated in the presence and absence of NAD^+^ and the observed differences in peptide deuteration were mapped onto the crystal structure of AbTir-TIR (Fig. 4A). Overall, the addition of NAD^+^ resulted in decreases in deuterium uptake by AbTir-TIR across multiple deuterium incubation time points (Fig. 4B). Notably, the deuterium uptake at the DD and EE loops located at the dimeric interface observed in X-ray structures 8G83 and 7UWG were significantly decreased at the earliest time point (10 sec). In addition, several AbTir-TIR peptides exhibited EX1 like exchange kinetics in the absence of NAD+. These regions are highlighted by cyan, red and purple bars in Fig 4B and color-coded accordingly in Fig 4C and D. The stacked isotopic envelops for representative peptides in these regions (Fig 4C) display progressive bimodal envelops characteristic of EX1 kinetic behavior in the unliganded form. When mapped onto the structures of AbTir-TIR, these areas, correspond to the amino-terminal A helix (FVRPLAETLQQL), the central B helix, the BC loop, the C β-strand (RQKIDSGLRNSKYGTVVL) and the carboxyl-terminal E helix (IAHQLAD) (Fig. 3, 4C, D and sFig 1) [26]. These peptides were found to colocalize, and the rate of formation of the high m/z species in the bimodal envelop was observed to be similar for all three regions. These observations reflect large, slow and cooperative conformational changes of AbTir-TIR in the absence of NAD^+^. Upon substrate binding, the progression of bimodal behavior is significantly reduced suggesting that NAD^+^ restricts or decreases the rate of these conformational changes. The NAD+ analog 3AD bound cryoEM structure (7UXU) exhibits a large conformational change in the BB loop and B Helix region as well as unique molecular assemblies in comparison to unliganded X-ray structures. Peptides exhibiting reduced deuterium uptake, large conformational changes and unique kinetics when mapped on the monomers of unliganded (8G83 and 7UWG) or ligand-bound form of AbTir (7UXU), with the exception of the B helix are not in direct contact with the substrate and instead map to molecular assembly TIR interfaces, loops, core β strands and helices (Fig 3 and Fig 4A [26, 44, 45].

**Figure 4.**
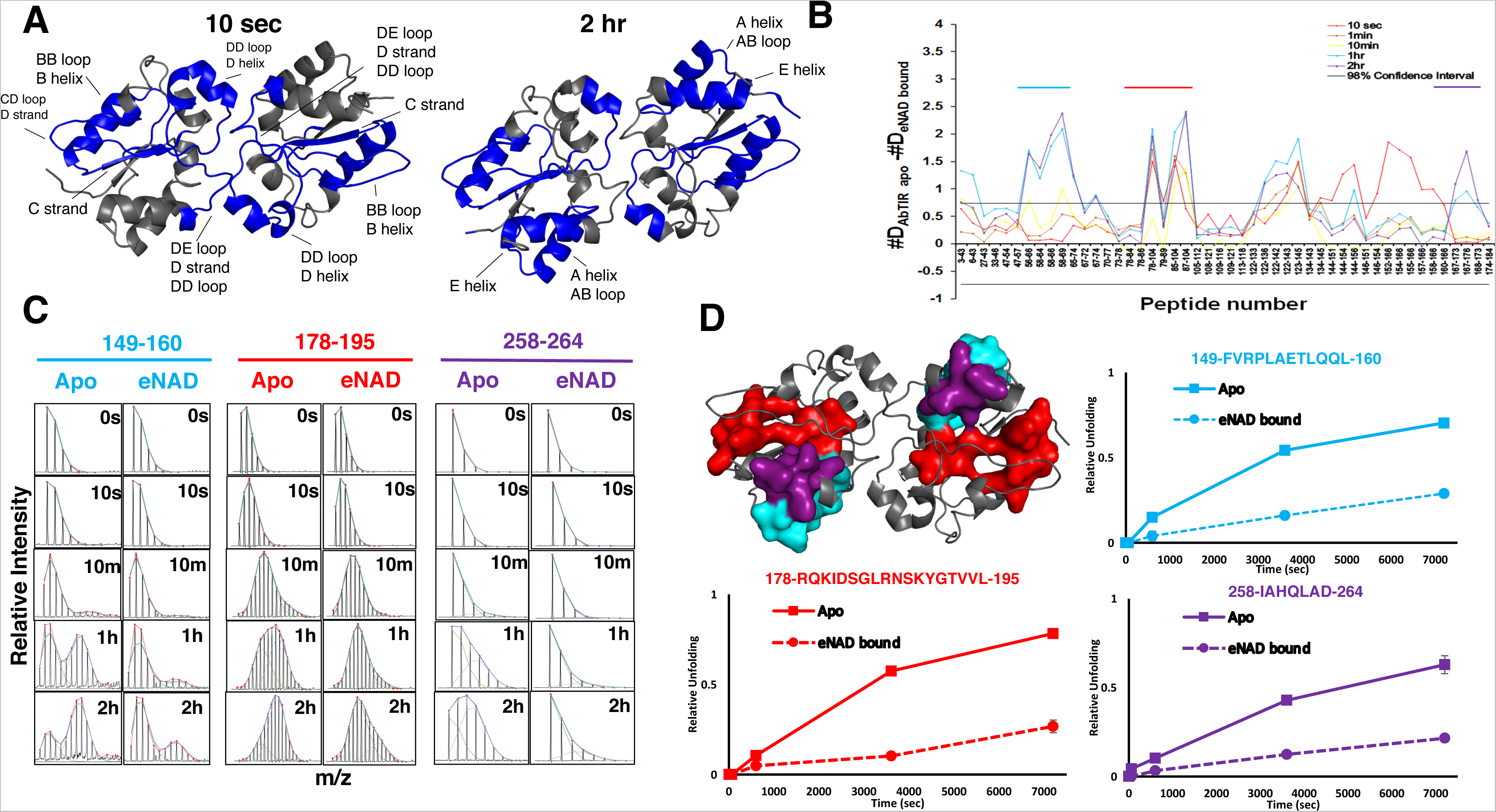
HDX-MS of AbTir-TIR^wt^ exhibit EX1 kinetics and conformations upon binding NAD+ NAD+ binding results in large decreases in deuterium uptake in AbTir-TIR^wt^. Significant EX1 exchange kinetics consistent with large conformational changes are observed for AbTir-TIR^wt^, in the absence of NAD and weakened upon ligand binding. A. Regions with significant decreases in deuterium uptake after 10sec and 2hr incubation are mapped on the structure of AbTir-TIR. B. The difference plot (Apo – εNAD bound) in absolute deuterium uptake at each incubation time point (color coded) and for each peptide (x-axis) C. Stacked spectra of representative spectra showing the signature bimodal isotopic envelops of EX1 kinetic regime of exchange. Apo and NAD bound states are shown. Peptides are color coded as in panel D. D. Surface and ribbon representation of AbTir-TIR^wt^. Surface representation shows peptides displaying bimodal EX1 behavior. Plots show the progressive accumulation of the high m/z species deconvolved from the bimodal isotopic envelop as a function of time of deuterium incubation for the apo (solid) and eNAD bound (dashed) AbTir-TIR^wt^.

## Discussion

Our structural studies of the *Acinetobacter baumannii* TIR domain (AbTir-TIR) are consistent with results reported by Manik et al. and reveal similarities and differences among bacterial, plant and mammalian TIR proteins [7, 8, 21, 22, 26]. A least-squares comparison with other bacterial TIR domain counterparts exhibits overall similarities of loops and interactions. Consistent with bacterial TIRs reported to date, the crystal structure of AbTir-TIR retains a similar overall core TIR domain fold with dimerization interactions as well as confirmed NAD+ hydrolase activity [22, 25].

NAD+ hydrolase activity of recombinant AbTir-TIR used for structural studies was confirmed by several different methods. LC/MS analysis indicates the ability of the recombinant AbTir-TIR^wt^ to hydrolyze NAD+ and produce novel cADPR products [11, 16, 17, 28]. Additionally, inactivated mutant AbTir-TIR^E208A^ has significantly reduced enzymatic activity. To assess the effect of AbTir-TIR expression directly in bacteria that produce them label-free live imaging using 2pFLIM was employed. 2pFLIM is a minimally invasive spectroscopic method used increasing in biological, biomedical and cancer applications [36, 37, 46–51]. 2p-FLIM uses the intrinsic fluorescence of NAD(P)H to monitor cellular function and biomedical applications including tissue morphology and high-density protein arrays [52–56]. Variances in both fluorescence intensity and fluorescence lifetimes are observed in bacteria expressing active (AbTir-TIR^wt^) and inactivated (AbTir-TIR^E208A^). Importantly, significant differences in the fluorescence lifetimes, which is independent of the protein concentration is observed among active and inactivated forms of AbTir-TIR. Differences among non-expressing (-IPTG) and expressing (+IPTG) active AbTIR^wt^ and inactive AbTIR^E208A^ fluorescence intensities and fluorescence lifetimes could be influenced by a number of factors including induction efficiency, lac I operon control, leaky expression, auto-induction, TIR overexpression related toxicity, NAD+ processing and salvage pathways or some other unaccounted phenomenon.

Changes in fluorescence lifetimes may reflect differences between bound and unbound free forms of NAD(P)H among active or inactivated AbTir-TIR and the effect this may have in the bacteria themselves [36, 37]. These current studies do not address observations resulting from bacterial TIR-induced toxicity, IPTG driven metabolic differences or leaky plasmid expression. Although, we have not fully characterized the molecular pathways resulting in AbTir-TIR induced NAD(P)H differences further studies using bacterial isolates and strains that naturally express or are bacterial TIR domain hydrolases may help bear this out. 2p-FLIM may prove useful as a minimally invasive live, label-free method for evaluating microbial infection in bacteria, plants and animals. Future studies characterizing bacterial isolates and strains that naturally express, are absent or have naturally induced TIR domain-containing hydrolase protein expression may better reflect biology.

Sequence comparison reveals conservation of the C-Helix WxxxE motif that is important for TcpB microtubule binding and includes the highly conserved Glu (E) residue which is critical for the enzymatic function of nearly all NAD+ hydrolases [11]. Without NAD+ bound this region including the highly conserved E(208) and Trp (W204) identified to be important for cADPR are observed facing away from the potential ligand binding region. Positioning of the Glu (E208) and Trp (W204) in are consistent with observations reported by Manik et al. HDX-MS substrate mediated conformational changes are also consistent with differences observed in X-ray structures and 3AD (NAD+ analog) bound cryoEM structures.

HDX-MS studies reveal that a large part of AbTir-TIR’s exchange behavior is governed by large conformational changes displaying EX1 kinetics which are altered upon ligand binding. These HDX-MS observations suggest that AbTir undergoes large conformational rearrangement and sampling, indicative of a heterogenous and complex conformational landscape in its native state. The WxxxE motif is not seen to be differentially protected from deuterium uptake by the addition of substrate and is not part of the region undergoing EX1 type conformational changes. Indeed, the conformational changes described by the HDX-MS are distinct in comparison with observations of X-ray and NAD analog bound cryoEM AbTIR assemblies (8G83, 7UWG and 7UXU). Substrate-induced conformational changes and kinetics observed in solution by HDX-MS at TIR interfaces, loop and secondary structure may be reflective of the transition from an unbound inactive state to an active substrate-bound assembly state.

Bioinformatics analysis of the AbTir-TIR domain identified regions that are highly conserved among bacterial and select human TIR domain-containing proteins [11, 25, 27, 40]. Directed targeting of regions and residues involved in mediating conformational change and substrate binding may provide insight into novel therapeutics to mitigate TIR-mediated pathogenesis. Indeed, the several NAD analogs as well as the anti-sepsis TIR-specific small molecule antagonist TAKEDA-242 which protects against lethal influenza binds specifically to the Cys747 located at position 2 within the Wx_C747_xxE motif of TLR-TIR4 C helix. Small molecule targeting of the WxxxE motif and surrounding region, as observed by TAK-242 protection against lethal influenza infection may provide protection against additional microbial infections or diseases. This cysteine is observed in several TIR-TIR interfaces involving C-Helix interactions with the WxxxE motif being highly conserved in select extracellular TLRs[57]. Additionally, functionally important residues found on the adjacent helices involved regulating the BB loop position may be potential regulator. Changes in neighbor secondary structure elements, interactions and the highly conserved “box2” residues may affect BB Loop position among different TIR domains.

Experimental procedures:

Protein expression, purification, analytical ultracentrifugation and crystallization.

The protein expression construct containing StrepTag-AbTir-TIR-HisTag within the pet30a+ vector was obtained from Dr. Kow Essuman [11]. The construct for mutant AbTir-TIR^E208A^ was created using PCR-based site-directed mutagenesis using the AbTir-TIR^wt^ construct as a template. Recombinant AbTir-TIR ^wt^ ^or^ ^E208A^ protein was expressed, purified, crystallized and data collected similar to previously described [21, 22]. After nickel purification AbTir-TIR was concentrated using an amicon 15kDa MWCO and further purified using size exclusion chromatography (SEC) using a S200 column in buffer containing 25mM HEPES pH7.2, 150mM NaCl. Initial crystals of AbTir-TIR^wt^ were identified from commercial screens using vapor diffusion method and further optimized by micro- and streak-seeding to yield long thin crystal clusters. Despite the success of seeding to reproduce crystals, attempts to further optimize crystals using various strategies including varying seed stock, batch or sparse matrix rescreening, liquid-liquid diffusion or macro seeding failed to markedly improve crystal size or singularity. Minor improvements in crystal length were observed intermittently and usually associated with reduced nucleation events. Crystallization of AbTir-TIR^wt^ with NAM or NAD+ substrate either by soaking or co-crystallization methods produced similar starburst-like crystals but these did not yield single crystal diffraction beyond ~4Å and no complete data sets were collected.

Analytical ultracentrifugation sedimentation equilibrium measurements on AbTir-TIR samples prepared at 12, 7, and 5μM were performed using a Beckman Coulter Optima XL-I centrifuge at rotor speeds of 23, 26, and 29K rpm in the same buffer used for size exclusion. Data were acquired at 280 nm after 6, 12 and 18 hours of centrifugation and the resulting nine scans were globally analyzed using Heteroanalysis (https://core.uconn.edu/resource/biophysics/#au-software) with single species and monomer-dimer model. Based on the distribution of the residuals and the square root of the variance of the fit, the latter model was more appropriate. Sedimentation velocity was performed on AbTir-TIR^wt^ prepared at 10μM at 45K rpm. Analysis of the scans acquired at 280 nm using DCDT^+^ yielded a single species with a sedimentation coefficient, s_20,w_, of 2.8S[32, 33].

### Structure determination

The structure of AbTir-TIR^wt^ was determined by molecular replacement using the TcpB TIR domain structure (4LQC) as a template model [22, 58]. An initial Cα model was built using auto-build (Phenix) followed by manual building in Coot [58, 59]. Iterative manual building of discontinuous loops and termini followed Phenix refinement resulting in the AbTir-TIR model.

### NAD+ hydrolase activity assays

The ability of AbTir-TIR protein to cleave NAD+ was confirmed by HPLC metabolite measurements, Etheno-NAD^+^ Assay, EnzyChrom™ NAD/NADH Assay (Bioassay systems) and one dimensional NMR. HPLC metabolite measurement was performed in the presence of NAD+ incubated for 30 and 60 minutes using recombinant purified AbTir-TIR^wt^ protein used for crystallization as described previously[11]. Etheno-NAD+ Assay was performed using Nicotinamide 1, N6-ethenoadenine dinucleotide NAD (εNAD) Sigma N263 incubated with recombinant AbTir-TIR protein at 50uM in 10 mM HEPES pH 7.5, 150 mM NaCl buffer with a reaction volume of 100ul. Fluorescence was read using a Molecular Devices Spectramax plus^TM^ plate reader (at Ex/Em = 310/410) [34]. NAD levels and hydrolase activity of recombinant AbTir-TIR^wt^ ^and^ ^E208A^ were measured using the EnzyChrom™ NAD/NADH Assay (Bioassay systems, Hayward, CA). AbTIR-TIR^wt^ ^and^ ^E208A^ were incubated with 5 μM NAD for 10-30 minutes. Means (N=6) + SEM, were analyzed by 2-way ANOVA to compare differences in protein type over time, followed by Tukey’s post hoc analysis; * indicates p < 0.0001. AbTir-TIR^wt^ reduces cellular levels of NAD+ upon recombinant expression in *E. coli*. AbTIR-TIR^wt^ ^and^ ^E208A^ was induced by IPTG and cells were harvested 0-6 hrs after induction. Means (N=4) + SEM, were analyzed by 2-way ANOVA to compare differences in protein type over time, followed by Sidak’s post hoc analysis. Letters indicate statistically significant difference (p < 0.001). Measurement of hydrolase activity of AbTir-TIR expressed in T7 cells was performed by measuring NAD+ levels in cells before and after expression of AbTir-TIR^wt^ ^and^ ^E208A^ using the EnzyChrom™ NAD/NADH Assay (Bioassay systems. Briefly, T7 cells transformed with pet30A+ vectors encoding for StrepTag-AbTir-TIR-^HisTagwt^ ^and^ ^E208A^ were grown to mid-log phase (Abs600 = 0.6-0.8), induced with 0.5mM IPTG and incubated for 0-6 h. Cells were normalized by OD600 measurements, homogenized in NAD extraction buffer (supplied) and measured for NAD+ levels according to the manufacturer’s instructions. NADase activity of recombinant, purified AbTir-TIR was further verified by 1D NMR spectra that followed NADase activities, a total volume of 330 ml of 1 mM NAD^+^ was used with 20 ml D_2_O. Reactions were followed for 30 minutes using 16 scans for a total of 30 spectra by adding 300 mM of either AbTir-TIR^WT^ or AbTir-TIR^E208A^. Data were collected using a Varian 900 spectrometer at the Rocky Mountain 900 facility.

### 2pFLIM

NAD hydrolase activity of bacteria containing AbTir-TIR^wt^ or AbTir-TIR^E208A^ plasmid was further analyzed by 2p-FLIM. Samples were grown and induced to express AbTir-TIR^wt^ or AbTir-TIR^E208A^ as described for protein expression and hydrolase activity experiments. Bacteria were measured by optical density at 600nm (OD_600_) and adjusted by dilution to the same concentration prior to imaging. A customized confocal microscope (based on ISS Q2 laser scanning nanoscope) with single-molecule detection sensitivity was used for performing 2p-FLIM. The excitation source is a pulsed femtosecond laser (Calmar, 780nm, 90 fs pulse width, and 50MHz repetition rate) equipped with an ISS excitation power control unit. An incident wavelength of 780 nm was used for exciting NADH in cells or tissue samples. The excitation light was reflected by a dichroic mirror to a high-numerical-aperture (NA) water objective (60X; NA, 1.2) and focused onto the sample. The fluorescence was collected by single photon counting avalanche photodiodes (SPAD) through a dichroic beam splitter, Chroma short pass (750SP), and 460/55 band-pass filters, thus eliminating the scattered excitation light and collecting fluorescence from the NADH in the region of interest. The imaging in the Q2 FCS/SMD setup was performed with Galvo-controlled mirrors with related electronics and optics controlled through the 3X-DAC control card. The software module in ISS VistaVision for data acquisition and processing and the time-correlated single photon counting (TCSPC) module from Becker & Hickl (SPC-150) facilitate FLIM measurements and analyses. Statistical analysis of data was performed using GraphPad prism 8.0 and Origin 2021 software.

### H/D Exchange Mass Spectrometry (HDX-MS)

The uptake of deuterium following exposure to deuterated water (^2^H_2_O) is monitored by LC-MS in a bottom-up approach. Undeuterated (control) samples were first acquired to obtain a sequence coverage map using an experimental workflow as follows: 2 μL of 50 μM AbTIR in 25 mM Tris, 150 mM NaCl pH 7.4 was diluted with 98 μL ice-cold quench containing 50 mM glycine pH 2.5, 1 M guanidine-HCl, resulting in a 100 μL reaction. The experimental workflow for the deuterated samples for H/D exchange reactions was as follows: 2 μL of apo 50 μM AbTIR or in complex with 10 mM eNAD was incubated in 48 μL D_2_O buffer containing 25 mM Tris, 150mM NaCl, pD 7.0 at 25°C. The resulting 50 μL deuteration reactions were quenched at various times (10 sec, 1 min, 10 min, 1 hr and 2h) with 100 μL ice-cold quench (100 mM glycine pH 2.5, 1.5 M Guanidine) prior to injection. Using a LEAP autosampler for high throughput acquisition, deuterated and un-deuterated (control) samples were injected into a Waters HDX system equipped with an Acquity M-class UPLC with in-line pepsin digestion. The resulting peptides were trapped on an Acquity UPLC BEH C18 peptide trap and separated using an Acquity UPLC BEH C18 column followed by injection into a Waters Synapt G2Si mass spectrometer with ion mobility capabilities. Resulting peptides were identified using the ProteinLynx Global Server 3.0.3 (PLGS) from Waters. H/D exchange reactions were performed by dilution into a ^2^H_2_O based buffer, pH 6.7 at 25°C. Samples at all deuterium incubation time points (including undeuterated controls and fully deuterated controls) were acquired in triplicate. The centroid mass shift of peptides identified by PLGS was tracked as a function of deuterium incubation times using DynamX 3.0 software (Waters).

### Data availability

The crystal structure of *Acinetobacter baumannii* Tir-TIR (AbTIR-TIR) was deposited in the PDB under the accession code 8G83. All other data is contained within the manuscript or will be shared upon request (Greg Snyder).

**Table I.** X-ray diffraction and data collection

|  |  |
| --- | --- |
| AbTIR<sup>wt</sup> (aa134-269) PDB code 8G83 |  |
| No. of crystals | 1 |
| Beamline | 17-ID-1 AMX, NSLS-II |
| Wavelength (Å) | 0.920072 |
| Detector | EIGER |
| Crystal to detector distance(mm) | 248.844 |
| Rotation range per image | 0.2, 0-180 |
| No. of images | 1-900 (truncated 200-800) |
| Space group | P21 21 21 |
| Unit-cell parameters(Å,°) | 44.20 76.19 97.33, 90, 90, 90 |
| Molecules per asymmetric unit | 2 |
| Resolution limits (Å) | 28.86 3.031 (3.139-3.031) |
| Total reflections | 13278 (1261) |
| No. of unique observations | 6668(639) |
| Completeness (%) | 98.01 (95.09) |
| Multiplicity | 2.0 (2.0) |
| $R_{\text{merge}}$ | 0.05218 (0.308) |
| $R_{\text{meas}}$ | 0.0738 (0.435) |
| $R_{\text{p.i.m.}}$ | 0.05218 (0.308) |
| Mean I/sigma I | 10.06 (2.32) |
| $R_{\text{work}}$ | 0.2276 (0.3156) |
| $R_{\text{free}}$ | 0.2743 (0.2529) |
| CC(work) | 0.952 (0.760) |
| CC(free) | 0.934 (0.908) |
| Number of non-hydrogen atoms | 2129 |
| macromolecules | 2122 |
| Protein residues | 277 |
| RMS(bonds) | 0.006 |
| RMS(angles) | 0.79 |
| Ramachandran favored (%) | 94.14 |
| Ramachandran allowed (%) | 5.86 |
| Ramachandran outliers (%) | 0.00 |
| Rotamer outliers (%) | 0.00 |
| Clashscore | 4.58 |
| Average B-factor | 56.44 |
| macromolecules | 56.46 |
| solvent | 50.00 |

## Author contributions

G.G., J.C., C.T., S.B., S.N., A. Scudder., T.O. expressed protein, designed mutants, performed experiments, analyzed data and prepared figures. E.K, Y.W. A.Soares. carried out X-ray experiments. Y.W expressed, purified, and grew protein crystals. A.Soares. assisted in X-ray data collection and processing. E.K. built and refined the X-ray structure. J.O.O performed HDX-MS experiments, analyzed data, prepared figures and wrote the paper. D.B. designed and supervised AUC experiments, analyzed data, wrote AUC methods and edited the manuscript. J.S.R and E.E designed and performed 1D NMR experiments including analyzing data and writing 1D NMR methods. K.E. J.M. AD. designed protein expression constructs performed HPLC experiments, analyzed, interpreted data and provided feedback. M.LD.S, K.R., D.D., G.A.S. conceived, designed, and supervised individuals carrying out investigations, analyzed data, prepared figures, and wrote the manuscript.

## Conflict of Interest

The research for this article was performed while Dorothy Beckett was employed at the University of Maryland. The opinions expressed in the article are the author’s own and do not reflect the view of the National Institutes of Health, the Department of Health and Human Services, or the United States Government.

The authors declare that they have no other conflicts of interest with the contents of this article.

## This article contains supporting information

**Supporting Figure 1 (sFig. 1).**
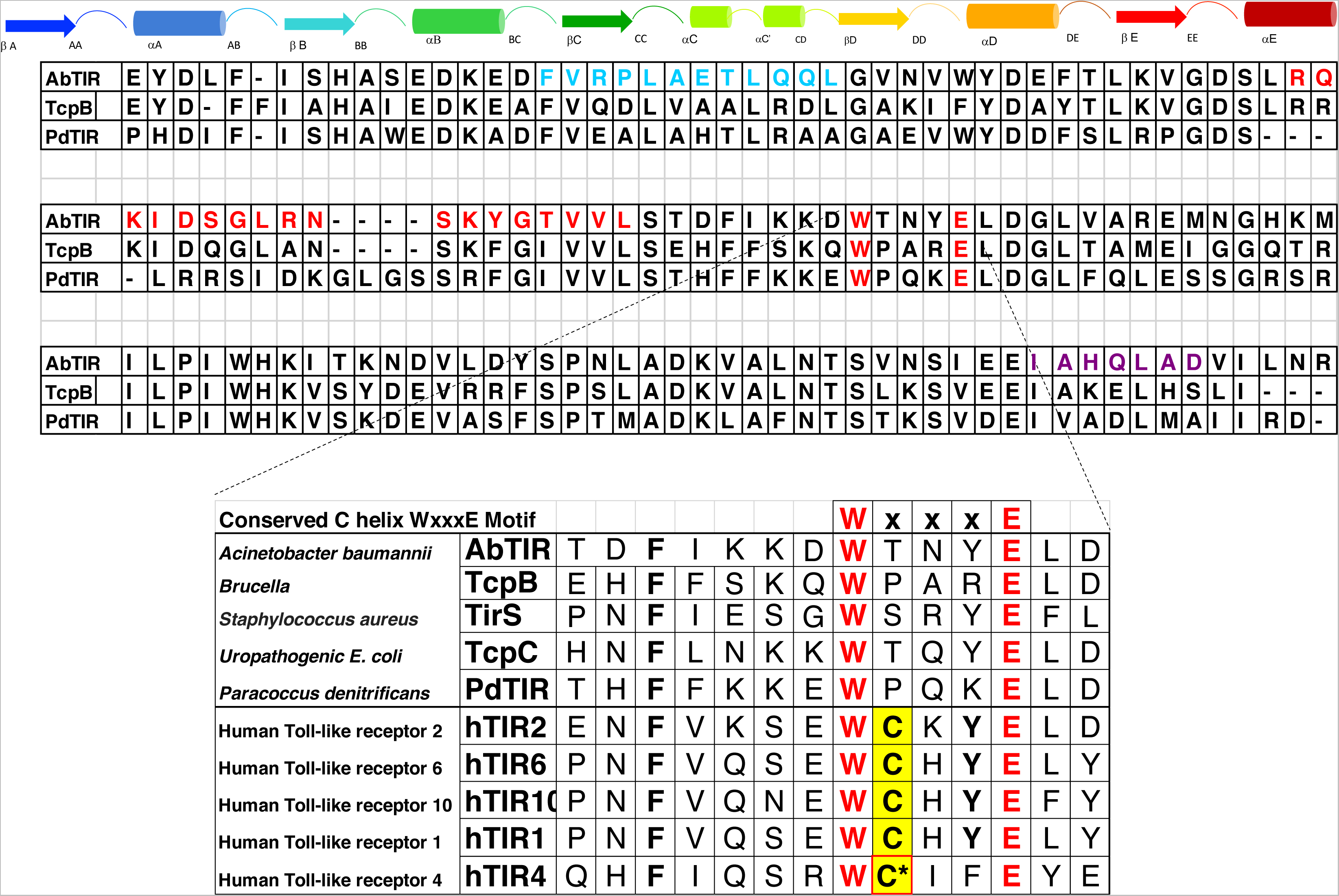
Sequence alignment of AbTir-TIR, TcpB, and PdTIR Sequence alignment of AbTir-TIR, TcpB, and PdTIR. The conserved WxxxE motif important for TcpB binding microtubule is highlighted in yellow. Inset shows conservation of bacterial WxxxE motif among bacterial and human TIR domain-containing proteins. Extracellular TLR-TIR proteins contain a cysteine within WxxxE motif and TLR4-TIR Cys747 is targeted by the small molecule inhibitor TAK-242 [57, 60].

**Supporting Figure 2 (sFig. 2).**
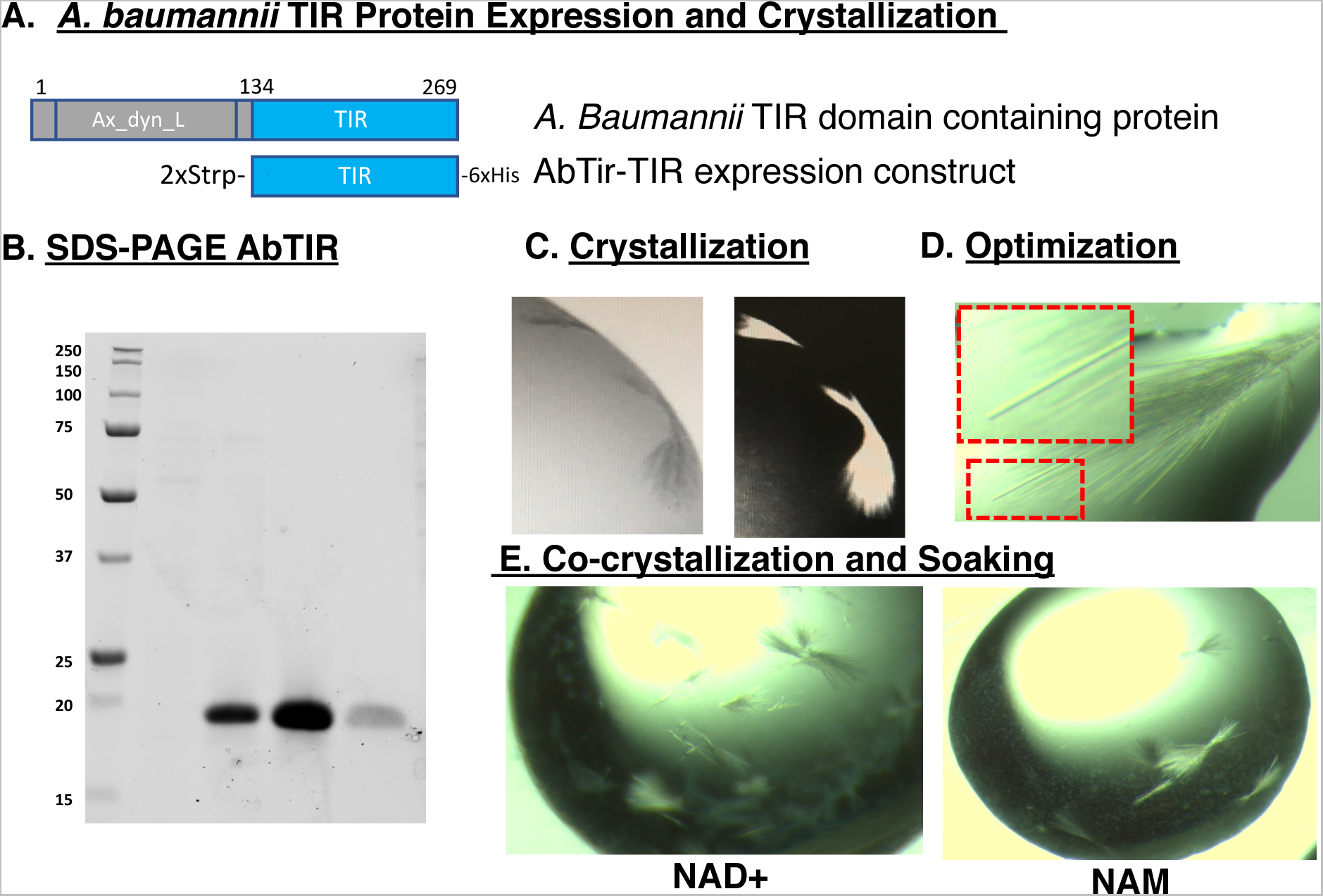
Acinetobacter baumannii TIR protein construct, expression, purification and crystallization. A. The TIR domain-containing protein from *A. baumannii*. The AbTir-TIR protein expression construct consists of the TIR domain portion, amino acids 134-269 (AbTIR) and amino 2 times (2x) streptavidin and carboxyl 6xHIS terminal purification tags. B. SDS-PAGE of Ni-NTA purified AbTir-TIRWT protein showing imidazole step gradient elution fractions. C. Crystals of AbTir-TIRwt. A. Initial broom-like crystals identified from commercial screening conditions MIDAS 2-19 (20 % v/v Jeffamine® M-2070, 20 % v/v Dimethyl sulfoxide) D. Streak seeding optimization of initial crystallization conditions. Inset box shows an example of thin rod like crystal used in diffraction and data collection experiments. E. Co-crystallization of AbTir-TIR in the presence of 1-5 mM NAD+ or NAM diffracted poorly.

**Supporting Figure 3 (sFig. 3).**
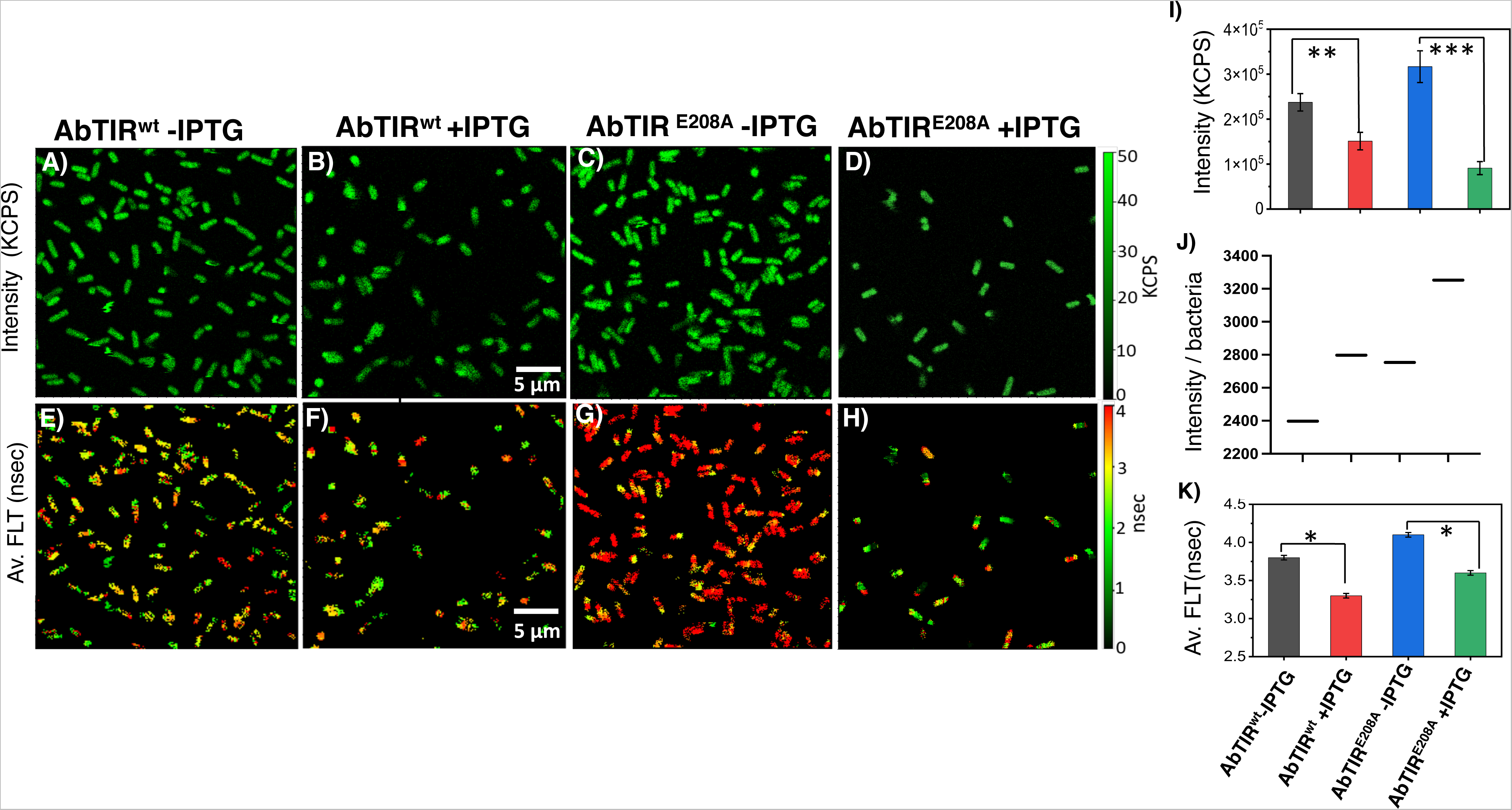
Live label-free 2pFLIM imaging of bacteria expressing AbTir-TIR. Two-photon fluorescence lifetime imaging microscopy (2p-FLIM) of live bacteria expressing either wild-type AbTir-TIRwt or inactivated mutant AbTir-TIR^E208A^. Panels A-D. The NADH fluorescence intensity images for bacteria uninduced (-IPTG) or induced (+IPTG) to express AbTir-TIRwt or E208Abacteria. Panels E-H. The lower panels depict images for the average fluorescence lifetimes. I. The top histogram shows the NADH average fluorescence intensity for panels A-D. J. The middle right histogram shows the average fluorescence intensity for panels A-D divided by the total number of bacteria imaged for each panel. K. The lower histogram shows the average fluorescence lifetimes for panels E-H. T-test showing P-value *≤ 0.05, ** ≤ 0.01 and *** ≤ 0.001.

**Supporting Figure 4 (sFig. 4).**
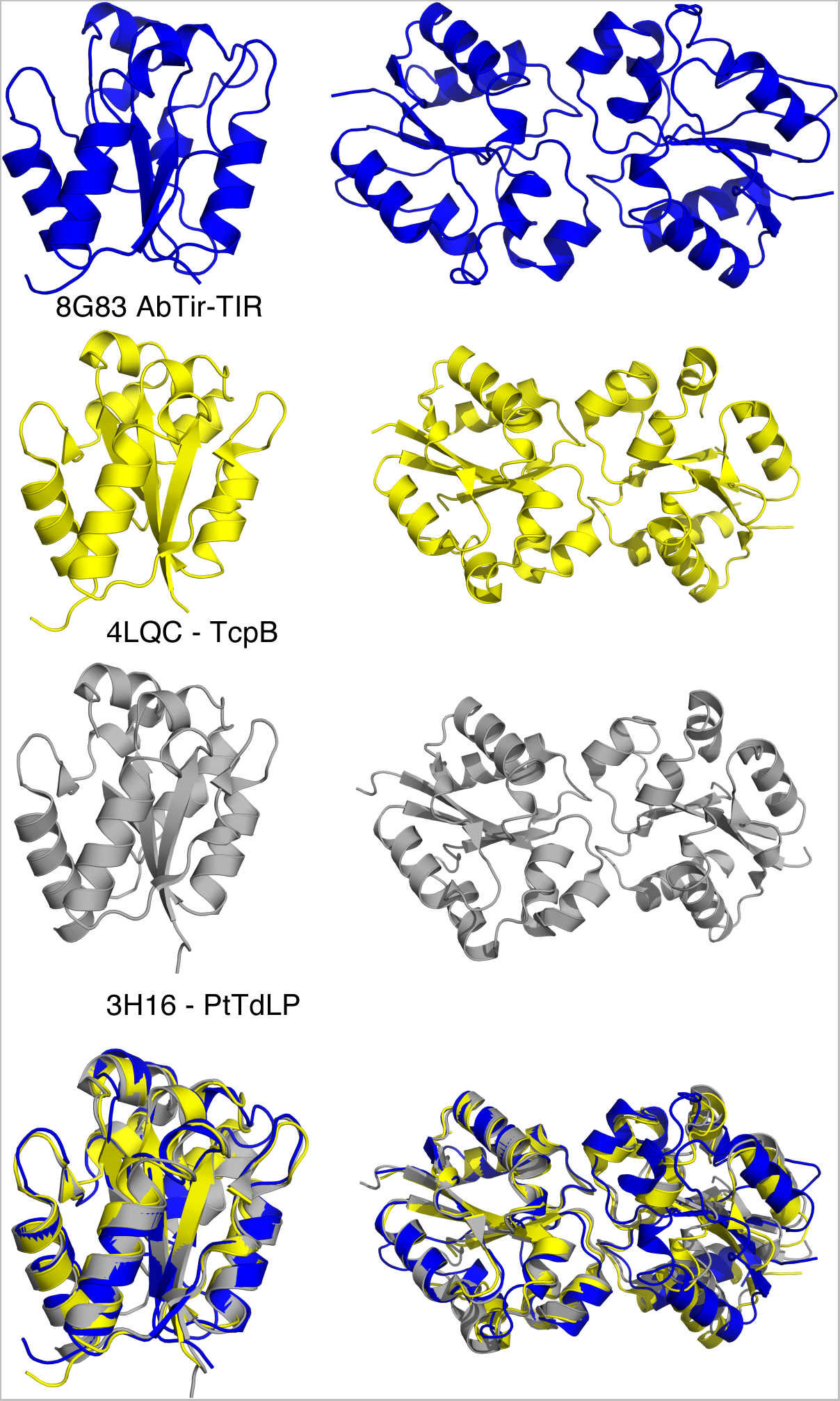
Comparison of AbTir-TIR with other bacterial TIR X-ray structures. AbTir-TIRwt monomer and dimer assemblies are similar to other unliganded bacterial TIR domain-containing protein structures. Ribbon diagram of crystal structures of AbTir-TIR (blue), Brucella spp. TcpB (yellow) and Paracoccus denitrificans PdTIR (grey). The left side depicts individual bacterial TIR domain monomers and the right side depicts bacterial TIR dimer as observed in the crystal. Superposition of unliganded AbTir-TIRwt, TcpB, and PdTIR show a high degree of similarity for overall loop positions and dimeric interactions.

**Supporting Figure 5 (sFig 5).**
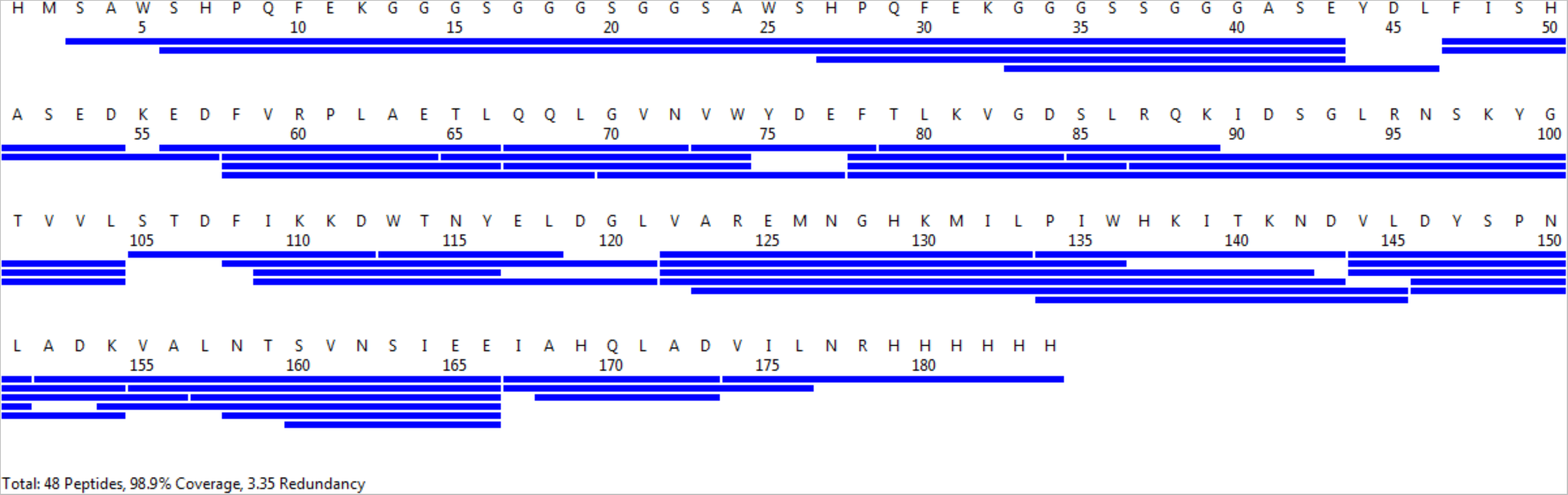
AbTir-TIR HDX-MS peptide coverage map. Hydrogen Deuterium Exchange Mass Spectrometry Peptide coverage map. Pepsin digestion of the AbTir-TIR TIR domain led to the detection of a total of 48 peptides covering 98.9% of the AbTir-TIR domain with 3.35 redundancy. The peptide coverage map was generated using MS Tools.

## Abbreviations

TIR: (Toll/Interleukin-1 receptor)
AbTir-TIR: (*Acinetobacter baumannii* TIR domain protein)
MDR: (multidrug-resistant)
HDX-MS: (hydrogen-deuterium exchange mass spectrometry)
cADPR: (cyclic ADP ribose)
PDB: (Protein Database)
FLT: (fluorescence lifetime)
2p-FLIM: (2-photon excitation with fluorescence lifetime imaging).

## Acknowledgments and Funding

The Center for BioMolecular Structure (CBMS) is primarily supported by the National Institutes of Health, National Institute of General Medical Sciences (NIGMS) through a Center Core P30 Grant (P30GM133893), and by the DOE Office of Biological and Environmental Research (KP1607011).

## Notes

### Competing Interest Statement

The authors have declared no competing interest.

## References

1. Turner, J.D., A bioinformatic approach to the identification of bacterial proteins interacting with Toll-interleukin 1 receptor-resistance (TIR) homology domains. FEMS Immunol Med Microbiol, 2003. 37(1): p. 13–21.

2. Zhang, Q., et al., TIR domain-containing adaptor SARM is a late addition to the ongoing microbe-host dialog. Developmental and comparative immunology, 2011. 35(4): p. 461–8.

3. Waldhuber, A., et al., A Comparative Analysis of the Mechanism of Toll-Like Receptor-Disruption by TIR-Containing Protein C from Uropathogenic Escherichia coli. Pathogens, 2016. 5(1).

4. Cirl, C., et al., Subversion of Toll-like receptor signaling by a unique family of bacterial Toll/interleukin-1 receptor domain-containing proteins. Nature medicine, 2008. 14(4): p. 399–406.

5. Radhakrishnan, G.K. and G.A. Splitter, Biochemical and functional analysis of TIR domain containing protein from Brucella melitensis. Biochemical and biophysical research communications, 2010. 397(1): p. 59–63.

6. Radhakrishnan, G.K., et al., Brucella TIR Domain-containing Protein Mimics Properties of the Toll-like Receptor Adaptor Protein TIRAP. The Journal of biological chemistry, 2009. 284(15): p. 9892–8.

7. Shi, Y., et al., Structural basis of SARM1 activation, substrate recognition, and inhibition by small molecules. Mol Cell, 2022.

8. Maruta, N., et al., Structural basis of NLR activation and innate immune signalling in plants. Immunogenetics, 2022. 74(1): p. 5–26.

9. DiAntonio, A., J. Milbrandt, and M.D. Figley, The SARM1 TIR NADase: Mechanistic Similarities to Bacterial Phage Defense and Toxin-Antitoxin Systems. Front Immunol, 2021. 12: p. 752898.

10. Weagley, J.S., et al., Products of gut microbial Toll/interleukin-1 receptor domain NADase activities in gnotobiotic mice and Bangladeshi children with malnutrition. Cell Rep, 2022. 39(4): p. 110738.

11. Essuman, K., et al., TIR Domain Proteins Are an Ancient Family of NAD(+)-Consuming Enzymes. Curr Biol, 2018. 28(3): p. 421–430 e4.

12. Essuman, K., et al., The SARM1 Toll/Interleukin-1 Receptor Domain Possesses Intrinsic NAD(+) Cleavage Activity that Promotes Pathological Axonal Degeneration. Neuron, 2017. 93(6): p. 1334–1343 e5.

13. Williamson, D.H., P. Lund, and H.A. Krebs, The redox state of free nicotinamide-adenine dinucleotide in the cytoplasm and mitochondria of rat liver. Biochem J, 1967. 103(2): p. 514–27.

14. Gerdts, J., et al., SARM1 activation triggers axon degeneration locally via NADLJ destruction. Science, 2015. 348(6233): p. 453–7.

15. Gerdts, J., et al., Sarm1-mediated axon degeneration requires both SAM and TIR interactions. J Neurosci, 2013. 33(33): p. 13569–80.

16. Bayless, A.M. and M.T. Nishimura, Enzymatic Functions for Toll/Interleukin-1 Receptor Domain Proteins in the Plant Immune System. Front Genet, 2020. 11: p. 539.

17. Wan, L., et al., TIR domains of plant immune receptors are NAD. Science, 2019. 365(6455): p. 799–803.

18. Patot, S., et al., The TIR Homologue Lies near Resistance Genes in Staphylococcus aureus, Coupling Modulation of Virulence and Antimicrobial Susceptibility. PLoS Pathog, 2017. 13(1): p. e1006092.

19. Askarian, F., et al., A Staphylococcus aureus TIR domain protein virulence factor blocks TLR2-mediated NF-kappaB signaling. J Innate Immun, 2014. 6(4): p. 485–98.

20. Imbert, P.R., et al., A. EMBO J, 2017. 36(13): p. 1869–1887.

21. Snyder, G.A., et al., Molecular mechanisms for the subversion of MyD88 signaling by TcpC from virulent uropathogenic Escherichia coli. Proceedings of the National Academy of Sciences of the United States of America, 2013.

22. Snyder, G.A., et al., Crystal structures of the Toll/Interleukin-1 receptor (TIR) domains from the Brucella protein TcpB and host adaptor TIRAP reveal mechanisms of molecular mimicry. J Biol Chem, 2014. 289(2): p. 669–79.

23. Li, Y., et al., Type I IFN operates pyroptosis and necroptosis during multidrug-resistant A. baumannii infection. Cell Death Differ, 2018. 25(7): p. 1304–1318.

24. Essuman, K., et al., TIR Domain Proteins Are an Ancient Family of NAD. Curr Biol, 2018. 28(3): p. 421–430.e4.

25. Sikowitz, M.D., et al., Reversal of the substrate specificity of CMP N-glycosidase to dCMP. Biochemistry, 2013. 52(23): p. 4037–47.

26. Manik, M.K., et al., Cyclic ADP ribose isomers: Production, chemical structures, and immune signaling. Science, 2022. 377(6614): p. eadc8969.

27. Felix, C., et al., The Brucella TIR domain containing proteins BtpA and BtpB have a structural WxxxE motif important for protection against microtubule depolymerisation. Cell Commun Signal, 2014. 12: p. 53.

28. Horsefield, S., et al., NAD. Science, 2019. 365(6455): p. 793–799.

29. Miller, M.S., et al., Getting the Most Out of Your Crystals: Data Collection at the New High-Flux, Microfocus MX Beamlines at NSLS-II. Molecules, 2019. 24(3).

30. Johnson, M.L., et al., Analysis of data from the analytical ultracentrifuge by nonlinear least-squares techniques. Biophys J, 1981. 36(3): p. 575–88.

31. *Heteroanalysis*. https://core.uconn.edu/resources/biophysics#au-software.

32. Stafford, W.F., Boundary analysis in sedimentation transport experiments: a procedure for obtaining sedimentation coefficient distributions using the time derivative of the concentration profile. Anal Biochem, 1992. 203(2): p. 295–301.

33. Philo, J.S., Improved methods for fitting sedimentation coefficient distributions derived by time-derivative techniques. Anal Biochem, 2006. 354(2): p. 238–46.

34. Schultz, M.B., et al., Assays for NAD. Methods Mol Biol, 2018. 1813: p. 77–90.

35. Lee, E., et al., Human and Bacterial Toll-Interleukin Receptor Domains Exhibit Distinct Dynamic Features and Functions. Molecules, 2022. 27(14).

36. Berezin, M.Y. and S. Achilefu, Fluorescence lifetime measurements and biological imaging. Chem Rev, 2010. 110(5): p. 2641–84.

37. Becker, W., Fluorescence lifetime imaging--techniques and applications. J Microsc, 2012. 247(2): p. 119–36.

38. Xu, Y., et al., Structural basis for signal transduction by the Toll/interleukin-1 receptor domains. Nature, 2000. 408(6808): p. 111–5.

39. Rock, F.L., et al., A family of human receptors structurally related to Drosophila Toll. Proc Natl Acad Sci U S A, 1998. 95(2): p. 588–93.

40. Toshchakov, V.Y. and A.F. Neuwald, A survey of TIR domain sequence and structure divergence. Immunogenetics, 2020. 72(3): p. 181–203.

41. Snyder, G.A. and E.J. Sundberg, Molecular Interactions in Interleukin and Toll-like Receptor Signaling Pathways. Current pharmaceutical design, 2013.

42. Chan, S.L., et al., Molecular mimicry in innate immunity: crystal structure of a bacterial TIR domain. The Journal of biological chemistry, 2009. 284(32): p. 21386–92.

43. Alaidarous, M., et al., Mechanism of Bacterial Interference with TLR4 Signaling by Brucella Toll/Interleukin-1 Receptor Domain-containing Protein TcpB. J Biol Chem, 2014. 289(2): p. 654–68.

44. Shi, Y., et al., Structural basis of SARM1 activation, substrate recognition, and inhibition by small molecules. Mol Cell, 2022. 82(9): p. 1643–1659.e10.

45. Sporny, M., et al., Structural basis for SARM1 inhibition and activation under energetic stress. Elife, 2020. 9.

46. Lakowicz, J.R., Principles of Fluorescence Spectroscopy, Chapter 22,. Third ed. 2006: Springer.

47. Ray, K. and J.R. Lakowicz, Metal-Enhanced Fluorescence Lifetime Imaging and Spectroscopy on a Modified SERS Substrate. J Phys Chem C Nanomater Interfaces, 2013. 117(30).

48. Fixler, D., T. Nayhoz, and K. Ray, Diffusion Reflection and Fluorescence Lifetime Imaging Microscopy Study of Fluorophore-Conjugated Gold Nanoparticles or Nanorods in Solid Phantoms. ACS Photonics, 2014. 1(9): p. 900–905.

49. Penjweini, R., et al., Single cell-based fluorescence lifetime imaging of intracellular oxygenation and metabolism. Redox Biol, 2020. 34: p. 101549.

50. Niehorster, T., et al., Multi-target spectrally resolved fluorescence lifetime imaging microscopy. Nat Methods, 2016. 13(3): p. 257–62.

51. Barnoy, E.A., et al., An ultra-sensitive dual-mode imaging system using metal-enhanced fluorescence in solid phantoms. Nano Res, 2015. 8(12): p. 3912–3921.

52. Sun, Y., et al., Fluorescence lifetime imaging microscopy for brain tumor image-guided surgery. J Biomed Opt, 2010. 15(5): p. 056022.

53. Coda, S., et al., Fluorescence lifetime spectroscopy of tissue autofluorescence in normal and diseased colon measured ex vivo using a fiber-optic probe. Biomed Opt Express, 2014. 5(2): p. 515–38.

54. Skala, M.C., et al., In vivo multiphoton microscopy of NADH and FAD redox states, fluorescence lifetimes, and cellular morphology in precancerous epithelia. Proc Natl Acad Sci U S A, 2007. 104(49): p. 19494–9.

55. Sarder, P., D. Maji, and S. Achilefu, Molecular probes for fluorescence lifetime imaging. Bioconjug Chem, 2015. 26(6): p. 963–74.

56. Sun, Y., et al., Monitoring protein interactions in living cells with fluorescence lifetime imaging microscopy. Methods Enzymol, 2012. 504: p. 371–91.

57. Shirey, K.A., Lai, Wendy., Brown, Lindsey J., Blanco, Jorge, B. C G., Robert.,, and Y. Wang, Vogel, Stefanie N., Snyder, Greg A.,, Select targeting of intracellular Toll-interleukin-1 receptor resistance domains for protection against influenza-induced disease. Innate Immunity, 2019. 10.1177/1753425919846281.

58. Adams, P.D., et al., PHENIX: a comprehensive Python-based system for macromolecular structure solution. Acta crystallographica. Section D, Biological crystallography, 2010. 66(Pt 2): p. 213–21.

59. Emsley, P. and K. Cowtan, Coot: model-building tools for molecular graphics. Acta crystallographica. Section D, Biological crystallography, 2004. 60(Pt 12 Pt 1): p. 2126–32.

60. Kawamoto, T., et al., TAK-242 selectively suppresses Toll-like receptor 4-signaling mediated by the intracellular domain. Eur J Pharmacol, 2008. 584(1): p. 40–8.

